# Impaired conscious access and abnormal attentional amplification in schizophrenia

**DOI:** 10.1101/186304

**Authors:** L Berkovitch, A Del Cul, M Maheu, S Dehaene

**Author notes:** denotes co-first authorship (L Berkovitch, A Del Cul). Corresponding author: Lucie Berkovitch, Cognitive Neuroimaging Unit CEA DRF/JOLIOT, INSERM, Université Paris-Sud, Université Paris-Saclay, DRF/JOLIOT/NEUROSPIN/UNICOG, Bât. 145 – Point Courrier 156, F-91191 Gif-sur-Yvette Cedex, FRANCE.

## Abstract

Previous research suggests that the conscious perception of a masked stimulus is impaired in schizophrenia, while unconscious bottom-up processing of the same stimulus, as assessed by subliminal priming, can be preserved. Here, we test this postulated dissociation between intact bottom-up and impaired top-down processing and evaluate its brain mechanisms using high-density recordings of event-related potentials. Sixteen patients with schizophrenia and sixteen controls were exposed to peripheral digits with various degrees of visibility, under conditions of either focused attention or distraction by another task. In the distraction condition, the brain activity evoked by masked digits was drastically reduced in both groups, but early bottom-up visual activation could still be detected and did not differ between patients and controls. By contrast, under focused top-down attention, a major impairment was observed: in patients, contrary to controls, the late non-linear ignition associated with the P3 component was reduced. Interestingly, the patients showed an essentially normal attentional amplification of the P1 and N2 components. These results suggest that some but not all top-down attentional amplification processes are impaired in schizophrenia, while bottom-up processing seems to be preserved.

## 1. Introduction

Schizophrenia is a serious psychiatric disorder that affects approximately ~1% of the population worldwide and causes positive symptoms, such as delusions and hallucinations, negative symptoms, including withdrawal from social interactions and daily life activities, cognitive impairments, and disorganization syndrome. Experimental studies of visual masking have reproducibly revealed an elevated threshold for the perception of masked visual stimuli in schizophrenia (Butler et al., 2003; Charles et al., 2017; Dehaene et al., 2003a; Del Cul et al., 2006; Green et al., 1999, 2011; Herzog et al., 2004; Herzog and Brand, 2015; Plomp et al., 2013). For instance, in classical masking experiments in which the target-mask duration is manipulated, patients with schizophrenia typically need a longer delay between the two, compared to controls, to consciously perceive the target (Charles et al., 2017; Del Cul et al., 2006). Similarly, patients are less likely to report that they perceived an unexpected event during inattentional blindness (Hanslmayr et al., 2013) and show an exaggerated attentional blink effect compared to controls, associated with a decreased P300 (Mathis et al., 2012).

Theoretical models of conscious processing suggest that the conscious perception of a stimulus involves the bottom-up propagation of sensory signals through the visual hierarchy, as well as top-down amplification by late and higher-level integrative processes (Dehaene et al., 2003b; Dehaene and Changeux, 2011). Many brain areas and networks continuously process sensory information in an unconscious manner, but conscious access is thought to start when top-down attention amplifies a given piece of information, allowing it to access a network of high-level brain regions broadly interconnected by long-range connections (Baars, 1993; Dehaene, 2011; Dehaene and Changeux, 2011; Lafuente and Romo, 2006). This so-called global neuronal workspace integrates the new incoming piece of evidence into the current conscious context, makes it available to multiple others brain processors and verbally reportable.

Conscious access, in the face of incoming sensory evidence, has been likened to a “decision to engage” the global workspace (Dehaene, 2008; Shadlen and Kiani, 2011). Borrowing from the diffusion model (Ratcliff, 1978) according to which decisions are made through a noisy process that accumulates information over time until sufficient information is obtained to initiate a response, it has been proposed that a non-conscious accumulation of sensory evidence precedes conscious access (Vorberg et al., 2003). According to that hypothesis, peripheral perceptual processors would accumulate noisy samples arising from the stimulus, and conscious access would correspond to a perceptual decision based on this accumulation (Dehaene, 2011; King and Dehaene, 2014). Both the amount of sensory evidence (e.g. the contrast of a stimulus) and the attentional resources would modulate the rate of accumulation of sensory information per unit of time, or drift rate, and thus the likelihood of consciously perceiving the stimulus. According to these theoretical models, an elevated consciousness threshold could thus result from both a bottom-up perceptual impairment and/or an insufficient top-down attentional amplification.

The increased sensibility to visual masking in schizophrenia was initially interpreted as indicating a bottom-up deficit, as other experimental results suggest low-level visual impairments in schizophrenia (Butler et al., 2003; Cadenhead et al., 1998; Green et al., 2011). Indeed, an impaired visual P1 to low spatial frequency stimuli was repeatedly observed in schizophrenic patients and attributed to a specific magnocellular visual pathway dysfunction (Butler et al., 2005, 2007; Javitt, 2009; Kim et al., 2006; Martínez et al., 2012). Moreover, schizophrenic patients exhibit deficits in the auditory P50, which is normally reduced for the second paired stimuli compared to the first, but insufficiently so in patients compared to controls (Javitt and Freedman, 2015), even if this effect may also be due to a dampened response to the first stimulus (Yee et al., 2010). Finally, patients also suffer from an abnormal prepulse inhibition of startle responses, a paradigm in which a weak sensory stimulus (the prepulse) inhibits the elicitation of the startle response caused by a sudden intense stimulus (Bolino et al., 1994; Braff et al., 1992).

However, observing a reduced activity of early ERP components is not sufficient to conclude in favor of a purely bottom-up impairment in schizophrenia. Similar findings could indeed also stem from impaired top-down attentional processes. This latter explanation is worth considering given the widely acknowledge modulatory effect that attention may have on early brain activation including the mismatch negativity (Kasai et al., 1999; Oades et al., 1997; Sauer et al., 2017), the P1 (Feng et al., 2012; Hillyard and Anllo-Vento, 1998; Luck and Ford, 1998; Wyart et al., 2012), and probably the P50 in healthy controls (Guterman et al., 1992) and schizophrenic patients (Yee et al., 2010). An additional argument suggesting that bottom-up processing may not be responsible for the patients’ elevated consciousness threshold in masking experiments comes from the observation that subliminal processing can be fully preserved in schizophrenia patients, as reported in a variety of paradigms with masked words (Dehaene et al., 2003a) or digits (Del Cul et al., 2006), subliminal error detection (Charles et al., 2017) and response inhibition (Huddy et al., 2009; for a review, see Berkovitch et al., 2017). This argument rests upon the idea that subliminal priming merely reflects the feed-forward propagation of sensory activation (Fahrenfort et al., 2008; Lamme and Roelfsema, 2000).

In summary, evidence for early visual processing deficits in schizophrenia is inconclusive and could be due either to an impairment of bottom-up processing, or to a lack of appropriate top-down attentional modulation as suggested by previous work (Dima et al., 2010; Fuller et al., 2006; Gold et al., 2007; Luck et al., 2006; Plomp et al., 2013).

Here we tested the hypothesis that bottom-up information processing is intact while top-down attentional amplification is deficient in schizophrenia by recording high-density electroencephalography (EEG) in a visual masking paradigm. We systematically and orthogonally manipulated a bottom-up factor (the delay between the mask and the target) and a top-down factor (whether the stimuli were attended or unattended). Our goal was twofold. First, we probed the brain mechanisms by which attention amplifies the processing of masked stimuli in healthy controls, therefore lowering down their threshold for access to conscious report. Second, we evaluated which of these mechanisms are impaired in schizophrenic patients. The hypothesis of intact bottom-up processing predicts that, once attention is withdrawn, early event related potentials (ERPs) should be equally reduced in both patients and controls, without any difference between these two groups. On the other hand, the difference between attended and unattended conditions, which provides a measure of attentional amplification, should reveal a deficiency of top-down amplification in schizophrenia, eventually resulting in a reduction or suppression of the global cortical ignition typically associated with conscious perception in normal subjects (Del Cul et al., 2007; Sergent et al.,2005).

The present research capitalizes upon a previous study in which we demonstrated that event-related potentials could be used to monitor the successive stages of processing of a masked stimulus (Del Cul et al., 2007). In this previous work, a digit target was presented for a brief fixed duration (14 ms), and followed - after a variable stimulus-onset-asynchrony (SOA) - by a mask consisting of surrounding letters. A fixed amount of sensory evidence was therefore initially injected while a variable amount of time was available to accumulate the evidence before the processing of the mask disrupted it. ERPs were used to monitor the successive stages of visual information processing associated with unconscious processing and conscious vision. Following the subtraction of mask-evoked brain activity, a series of distinct stages were observed. First, the PI and the Nl components were shown to vary little with SOA, reflecting the unconscious processing of the incoming digits. Second, an intermediate negative waveform component (N2J linearly increased with SOA but stopped at a fixed latency with respect to the mask, suggesting an accumulation of evidence in occipito-temporal cortical areas and its interruption by the mask. Finally, the late P3 component showed a sigmoidal variation with SOA, tightly parallel to subjective reports of target visibility, thus suggesting that the P3 indexes an all-or-none stage of conscious access to perceptual information (see also e.g. Sergent et al., 2005).

In the present study, we aimed at replicating those findings as well as probing which of these stages persist when the very same stimulus (a masked digit) is presented under conditions of inattention (see Figure 1). By doing so, we intended to explore the interaction between the amount of masking (as modulated by target-mask SOA) and the availability of attentional resources, and to manipulate those variables while comparing schizophrenic patients and controls. In the focused attention condition, subjects were asked to focus their attention to the peripheral masked digits and to report their visibility (as in the original study by Del Cul et al., 2007). In the unattended condition, we maximized the withdrawal of attention from our masked stimuli through the use of a highly demanding concurrent task: subjects were asked to focus on small color changes presented at fixation and to report which color was predominant, while the same masked digits were presented in the periphery of the visual field. Because the digits were entirely task-irrelevant, presented at a parafoveal location and asynchronous with the color changes, all kinds of attention were withdrawn (executive attention, i.e. linked to the task; spatial attention, i.e. linked to the location of the stimulus; and temporal attention, i.e. linked to the timing at which the stimulus appears).

Based on our hypothesis of preserved feedforward and impaired top-down processing in schizophrenia, we predicted that, under inattention, the early sensory components indexed by PI, Nl and even N2 would remain present (though reduced by inattention) and identical in patients and controls. We also expected that attention would amplify these sensory components in order to facilitate the accumulation of sensory evidence from the masked stimulus, and that this amplification would be impaired in schizophrenia patients.

## 2. Material and methods

### 2.1. Participants

Sixteen patients with schizophrenia (mean age 37 years, range 25-51; 5 women) participated to the study. All were native French speakers. Patients met DSM-IV criteria (or schizophrenia or schizoaffective disorders and were recruited from the psychiatric department of Creteil University Hospital (Assistance Publique, Hôpitaux de Paris). They had a chronic course and were stable at the time of the experiment. A French translation of the Signs and Symptoms of Psychotic Illness Scale (SSPI) (Liddle et al., 2002) was used to evaluate their symptomatology, and chlorpromazine equivalents were calculated to assess whether there was significant correlations between symptoms, treatment and behavioural results.

The comparison group consisted of sixteen control subjects (mean age 35.5 years, range 21-51, 4 women). Comparison subjects were excluded for history of any psychotic disorder, bipolar disorder, recurrent depression, schizotypal or paranoid personality disorder. Patients and controls with a history of brain injury, epilepsy, alcohol or substance abuse, or any other neurological or ophthalmologic disorders were also excluded. Patients and controls did not differ significantly in sex, age and level of education (see Table 1). All experiments were approved by the French regional ethical committee for biomedical research (Hôpital de la Pitié Salpêtrière), and subjects gave written informed consent.

**Table 1.** Characteristics of participants.

| **Characteristics** | **Schizophrenic**<br>**mean (± s.d.)** | **Control**<br>**mean (± s.d.)** | **Statistical test**<br>**(test value, p-value, BF)** |
| --- | --- | --- | --- |
| Sample size | 16 | 16 | — |
| Age (y.o.) | 37.44 (±7.4) | 35.5 (±10.5) | $t_{26.99} = 0.60$<br>$p = 0.55$<br>$BF = 1/2.59$ |
| Gender (M/F) | 11/5 | 12/4 | $\chi^2_1 = 0.16$<br>$p = 0.69$<br>$BF = 1/2.75$ |
| Years of education (from first year of high school) | 7.9 (±2) | 8.9 (±3.3) | $t_{24.90} = -1.04$<br>$p = 0.31$<br>$BF = 1/1.97$ |
| SSPI* scale total score | 12.2 (±6.8) | — | — |
| Antipsychotic equivalence dose (CPZ-Eq., in mg) | 650.2 (±376.3) | — | — |

\* Sign and Symptom of Psychotic Illness.

### 2.2 Design and procedure

The experimental paradigm is summarized in Figure 1. We used a variant of the masking paradigm designed in our previous studies in normal and clinical populations (Charles et al., 2017; Del Cul et al., 2006. 2007). A target digit (1, 4, 6 or 9) was presented for a fixed duration of ~ 14 ms at a randomly chosen position among four (1.4 degrees above or below and 1.4 degrees right or left of the fixation cross). After a variable delay (stimulus onset asynchrony or SOA), a metacontrast mask appeared at the target location for 250 ms. The mask was composed of four letters (two horizontally aligned M and two vertically aligned E) surrounding the target stimulus location without superimposing or touching it. Four visibility levels (SOAs 27, 54, 80 and 160 ms) and a mask-only condition were randomly intermixed across trials. In the mask-only condition, the target number was replaced by a blank screen with the same duration (i.e. 14 ms). The fixation cross was surrounded by 5,6 or 7 successive colored circles which could be either blue or yellow. The presentation of each of these circles lasted for 100 ms, and the inter-stimulus interval between them was 413 ms (SOA = 513 ms).

The same exact sequence of stimuli was presented under two distinct conditions, which differed only in the requested task. Under the attended condition, subjects were asked to pay attention to the masked digits and give two behavioural responses: (1) decide whether the digit was larger or smaller than 5 (which provided an objective measure of target perception) and (2) report the digit visibility using a categorical response “seen” or “not seen” (which provided a subjective measure of conscious access). Under the unattended condition, participants had to estimate the predominant color of the rapid sequence of colored circles surrounding the fixation cross. Note that the peripheral stimuli always appeared between the 2^nd^ and the 3^rd^ colored cirdes, while participants were still forced to pay attention to the central task because not enough evidence was yet delivered to accurately decide which of the 2 colors was the most frequent (given that the number of circles varied between five and seven). On each trial, feedback informed the subjects whether their answer was correct or not in order to reinforce their motivation and help them to maintain attention. At the end of the unattended blocks, participants were asked whether they noticed anything in their peripheral visual field.

Instructions for both attended and unattended tasks were given at the beginning of the experiment and were repeated before each block (attended or unattended). Subjects were asked to complete four blocks of trials: two “attended” blocks (A) and two “unattended” blocks (U), in A-U-U-A order for half of the subjects and in U-A-A-U order for the other half. There were 640 trials in total (320 unattended and 320 attended), i.e. 64 trials in each combination of attention (2 levels) and masking (5 levels, i.e. SOA = 27,54,80, or 160 ms, plus the mask-only condition).

**Figure 1.**
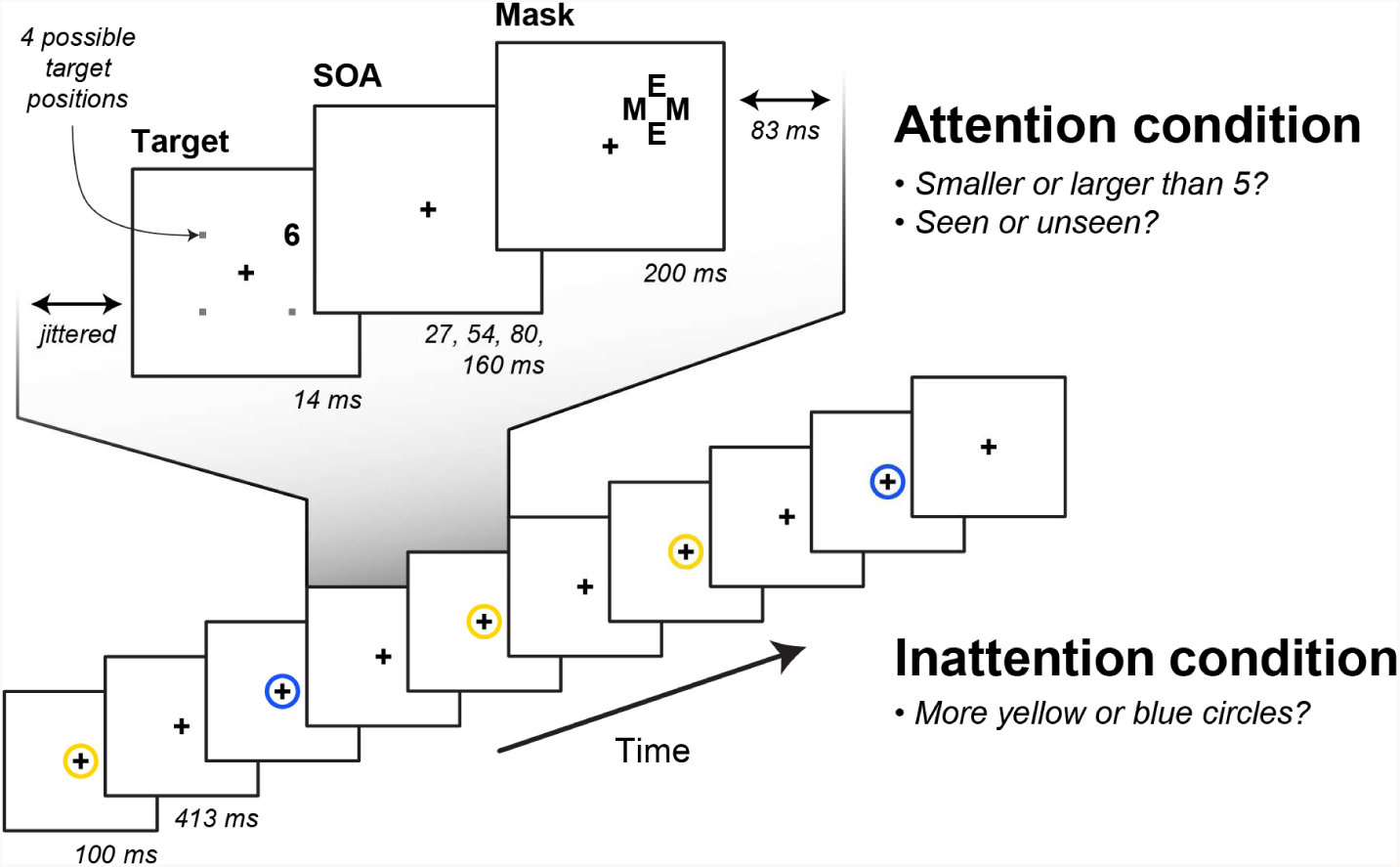
Experimental design. On each trial, subjects viewed a stream of small circles presented at fixation, with a brief presentation of a masked digit at one of four possible locations in the periphery of the visual field. The same exact sequence of stimuli was presented in two distinct experimental conditions. In the attended condition, subjects were asked to compare the target digit to a fixed reference of 5 (two alternatives forced-choice, objective task), then report whether they could see it or not (subjective visibility task). Ihe delay between the target and the metacontrast mask (SOA) varied between 27 and 160 ms in older to modulate the amount of masking, in the unattended condition, subjects had to estimate the predominant color of small circles surrounding the fixation cross, thus withdrawing attention from the irrelevant peripheral digit.

### 2.3. Behavioural data analysis

For each subject, several behavioural parameters were measured separately in each SOA condition. In the attended condition, we measured the performance in comparing the target against 5 (objective measure of conscious access) and the rate of seen trials (subjective measure of conscious access). In the unattended condition, we measured the performance in estimating which color was more frequent. Analyses of variance (ANOVAs) were conducted on each of those behavioural measures, with SOA as a within-subject factor and group (patients or controls) as a between-subject factor. Within the patient group, Pearson correlation coefficients were computed between behavioural measures and variables such as the clinical scale (SSPI scale, measuring the extent of positive, negative, and disorganisation symptoms, Liddle et al., 2002) and antipsychotic treatment posology (chlorpromazine equivalent). A measure of sensitivity (d’) was computed by confronting subjective visibility (seen versus not seen) against the presence or absence of a target (target versus mask-only trials).

### 2.4. ERP methods

EEG activity was acquired using a 128-electrode geodesic sensor net referenced to the vertex, with an acquisition sampling rate set to 250 Hz. We rejected voltage exceeding ± 200 μV. transients exceeding ±100μV, or electro-oculogram activity exceeding ±70μV (IV. The remaining trials were averaged in synchrony with mask onset, digitally transformed to an average reference, band-pass filtered (0.5 - 20 Hz) and corrected (or baseline over a 250 ms window during fixation at the beginning of the trial. Contralateral activity is represented conventionally on the left hemisphere and ipsilateral activity on the right one. The activity observed on mask-only trials was subtracted from that on trials in which the target was effectively presented, thus isolating the target-evoked activity.

In order to quantify the modulatory effect of SOA on EEG activity, linear regression models were fitted at the subject-level on the trial-averaged EEG signals, separately at each electrode and each time-point using the values of SOA as a parametric modulator (combined with an offset variable) of the EEG response. Group averaged regression coefficients (beta) corresponding to SOA were estimated, and *R*^2^ values (i.e. proportion of explained variance) are reported as an unbiased and normalized measure of the quality of fit.

ERP components were identified based on latencies, topographical responses (contralateral PI and Nl, bilateral N2 and P3) and previous work (Del Cul et al., 2007). For each subject, under each SOA and attention condition and for each digit-evoked ERP component, the EEG signals were averaged over corresponding clusters of electrodes and time windows (P1: 65-110 ms over parietotemporal electrodes: Nl: 125-200 ms over parietotemporal electrodes; N2: 200-300 ms over fronto-central electrodes; P3: 300-500 ms over fronto-central electrodes; see Del Cul et al., 2007).

In order to assess effect of experimental variables, we conducted analyses of variance (ANOVAs) separately for each these ERP components on their corresponding averaged amplitude (over electrodes and time points) with SOA (categorically recoded) and attention condition (attended or not) as within-subject factor and group (patients or controls) as a between-subject factor. We also compared the amplitude of each component against zero using a *t*-test in order to identify which of these components significantly persisted in the unattended condition.

### 2.5. Source localization

Cortical current density mapping was obtained using a distributed model consisting of 10.000 current dipoles. Dipole locations were constrained to the cortical mantle of a generic brain model built from the standard brain of the Montreal Neurological Institute, and warped to the standard geometry of the EEG sensor net. The warping procedure and all subsequent source analysis and surface visualization were performed using Brainstorm software (http://neuroimage.usc.edu/brainstorm) (Tadel et al., 2011). EEG forward modelling was computed with an extension of the overlapping.spheres analytical model (Huang et al., 1999). Cortical current maps were computed from the EEG time series using a linear inverse estimator (weighted minimum-norm current estimate or wMNE: see Baillet et al., 2001, for a review). We localized the sources separately for each subject and computed a group average that was then smoothed at 3 mm FWHM (corresponding to 2.104 edges on average), and thresholded at 40% of the maximum amplitude (cortex smoothed at 30%).

### 2.6. Statistical comparisons

Because many of the hypotheses at stake lie on an absence of difference (e.g. preserved feedforward processing in schizophrenic patients), besides frequentist statistics (values of the statistic, e.g. *ts* or *Fs*, as well as *p*-values are reported), we also conducted Bayesian statistics whenever required. Contrary to frequentist statistics, Bayesian statistics symmetrically quantify the evidence in favour of the null (H_0_) and the alternative (H_1_) hypotheses, therefore allowing to conclude in favour of an absence of difference (Wagenmakers et al., 2010). To do so, the *BayesFactor* package (http://bayesfactorpcl.r-forge.r-project.org) implemented in R (http://www.r-project.org) was used. Bayes Factor were estimated using a scale factor of r = 0.707. For each Bayesian statistical test, the corresponding Bayes factor (BF_10_ = p(data|H_1_)/p(data| H_0_)) is reported. Even though threshold values of Bayes factors have been proposed (e.g. a BF larger than 3 is usually taken has providing substantial evidence), a BF value of *x* can directly be interpreted as the observed data being approximately *x* times more probable under the alternative compared to the null hypothesis. When BFs favored the null hypotheses (i.e. BF_10_ < 1), we directly reported the inverse Bayes factor (i.e. BF_01_ = 1/BF_10_) quantifying the evidence in favor of the null compared to the alternative hypothesis.

## 3. Results

### 3.1. Behaviour

Behavioural results appear in Figure 2. As concerns the main digit-related task, under the attended condition, a main effect of SOA was observed on both objective performance (F_1,30_ = 184.02, p < 0.001)and subjective visibility (F_1,30_ = 287.17, p < 0.001).

Objective performance was significantly lower for patients compared to controls (73.7% vs. 80.7%. group effect F_1,30_ = 7.44, p = 0.011), but a significant group × SOA interaction (F_3,90_ = 3.14, p = 0.029) reflected the fact that this difference was significant only at the longest SOAs, i.e. 80 ms and 160 ms (F_1,30_ = 11.21, p = 0.002), not at the shortest SOAs 27 ms and 54 ms (F_1,30_ = 2.78, p = 0.110, BF = 1/1.8). Importantly, objective performance remained higher than chance in both groups (controls: 66.2%, t_31_ = 6.19, p < 0.001, patients: 61.7%, t_31_ = 5.624, p < 0.001).

Subjective visibility was also affected by a group × SOA interaction (F_3,90_ = 5.83, p = 0.001). Indeed, patients reported a significantly lower visibility at SOAs 80 ms and 160 ms (patients: 81.1% vs. controls: 91.3%; F_1,30)_ = 4.53, p = 0.042), and a significantly higher visibility in the mask-only and the 27 ms SOA conditions (14.3% vs. 3.9%, F_1,30_ = 5.53, p = 0.026) compared to controls. No difference was observed between the two groups at SOA 54 ms (F_1,30_ = 0.083, p = 0.780, BF = 1/2.9). Measures of sensitivity (d’) confirmed that patients were less able than controls to detect the target digit when SOAs were long (80 ms: t_27.7_ = −2.66, p = 0.013; 160 ms: t_17.6_ = −2.55, p = 0.020), while no significant difference was observed for short SOAs (27 ms: t_27.3_ = 1.44, p = 0.162, BF = 1/1.4; 54 ms: t_29.9_ = −1.03, p = 0.312, BF = 1/2.0).

Objective and subjective visibility were strongly correlated within subjects in both groups, and the strength of this correlation did not significantly differ between the two groups (mean Pearson r for controls: 0.97 vs. 0.96 foe patients, t_29.85_ = 0.30, p = 0.764, BF = 1/2.9). However, the patients’ objective performance was neither significantly correlated with the treatment(Pearson r = 0.095, t_14_ = 0.36, p = 0.725, BF = 1/5.0), nor with the clinical score (Pearson r = −0.28, t_14_ = −1.07, p = 0.304, BF = 1/3.1). Subjective performance showed a weak trend towards a negative correlation with treatment (across all SOAs: Pearson r = −0.50, t_14_ = −2.18, p = 0.046, BF = 1.4, for SOAs = 80 or 160 ms: Pearson r = −0.47, t_14_ = −1.99, p = 0.066, BF =1.0), but this correlation was strongly driven by one participant’s results (chlorpromazine equivalent: 1550 mg per day, subjective visibility across all SOAs: 16.0%; correlation after excluding the participant: Pearson r = −0.16, t_13_ = −0.59, p = 0.567, BF = 1/4.4). Finally, the clinical score was not correlated with subjective visibility (all SOAs: Pearson r = 0.00, t_14_ = 0.00, p = 0.997, BF = 1/5.3; for SOAs = 80 or 160 ms: Pearson r = −0.14, t_14_ = −0.54, p = 0.596, BF = 1/4.6).

As concerns the distracting task, under the unattended condition, performance in the central color task was lower for patients compared to controls (81.9 % vs. 90.9 %, F_1,30_ = 11.48, p = 0.002) There was no main effect of SOA (F_4,120_ = 0.39, p = 0.817, BF = 1/43.0) nor a group × SOA interaction (F_4,120_ = 1.16, p = 0.331, BF = 1/13.3). Within the patient group, performance was neither significantly correlated with treatment (Pearson r = 0.43, t_14_ = 1.791, p = 0.095, BF = 1/1.3) nor with clinical score (Pearson r = −0.45, t_14_ = −1.91, p = 0.077, BF = 1.1).

After the experiment, all subjects reported that they noticed the presence of the peripheral masked stimuli in the unattended condition, but that these stimuli could not be precisely identified and did not prevent them from estimating the dominant color of the central circles.

**Figure 2.**
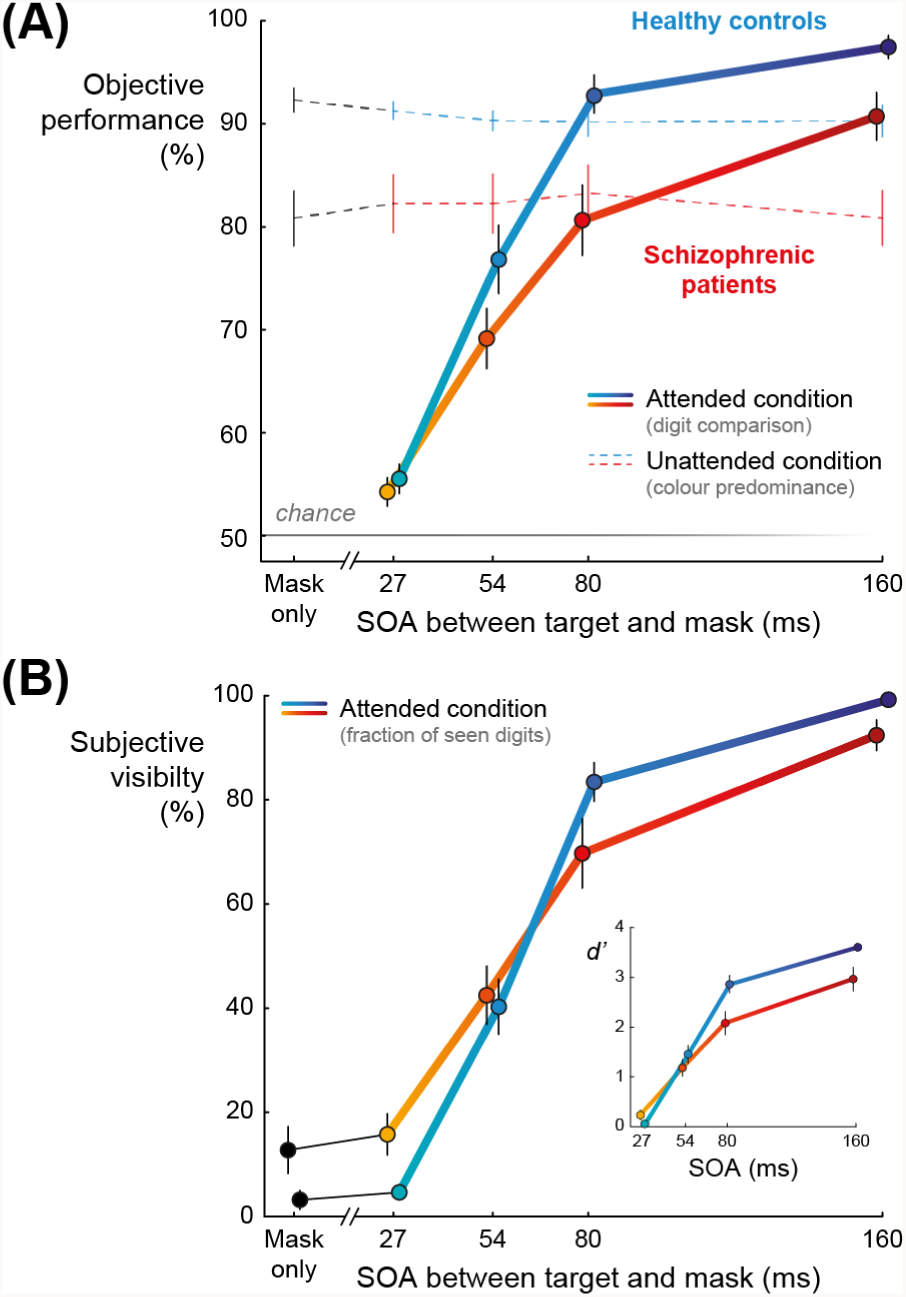
Behavioural results. (A) Objective performance as a function of SOA in the attended (comparing the masked digit to 5, solid lines) and the unattended conditions (estimating the predominant color of small circles surrounding the fixation cross, dashed lines). Error bars represent one standard error of the mean. Healthy controls (blue lines) performed better than schizophrenic patients (red lines) in both conditions. There was no effect of SOA in the unattended condition. (B) Subjective visibility of the masked digit and d’ measures as a function of SOA in the attended condition. Error bars represent one standard error of the mean. Healthy controls (blue lines) reported higher visibility and had higher d’ than schizophrenic patients (red lines) for long SOAs (i.e. 80 and 160 ms). Schizophrenic patients reported higher visibility than controls in the mask-only and the 27 ms SOA conditions but d’ measures did not significantly differ between the two groups for short SOAs (i.e. 27 and 54 ms).

### 3.2. EEG activity evoked by the target

Target-evoked brain activity is shown in Figure 3A in the case of the longest SOA (i.e. 160 ms) in the attended condition for both groups. At least five different components specific to conscious EEG visual responses could be identified: contralateral P1 (peaking at 88 ms post-target) and N1 (160 ms) followed by bilateral N2 (252 ms), P3a (324 ms) and P3b (392 ms). Scalp topographies and corresponding sources reconstruction are shown at specific time points (0, 88, 160, 252, 324, 392 and 600 ms).

First, at 88 ms and 160 ms (corresponding respectively to P1 and N1 components), brain activity elicited by the target was restricted to contralateral occipito-temporal regions (conventionally displayed on the left hemisphere) in both groups, reflecting the activation of early visual areas. The activity was slightly more diffuse and ventral in the patient group at 160 ms. At 252 ms (with a topography corresponding to the N2/P3a component), the activity spread to the ipsilateral hemisphere and moved forward in the postero-lateral part of the inferior temporal gyrus, including the visual number form area (Shum et al., 2013), and anterior prefrontal activity was detected. Then, at 324 ms, as a posterior P3b began to emerge in the scalp topography, the source activity spread bilaterally into the ventral stream, though more pronounced in the contralateral hemisphere, as well as in the inferior prefrontal and parietal cortices. Finally, at 392 ms (corresponding to the full-blown P3b component), activity became intense and fully bilateral in both groups, reaching ventral and dorsolateral prefrontal as well as parietal regions, especially in the control group. At 600 ms, in both groups, activity strongly decreased in the occipital lobes while remaining sustained in anterior frontal and temporal regions.

**Figure 3.**
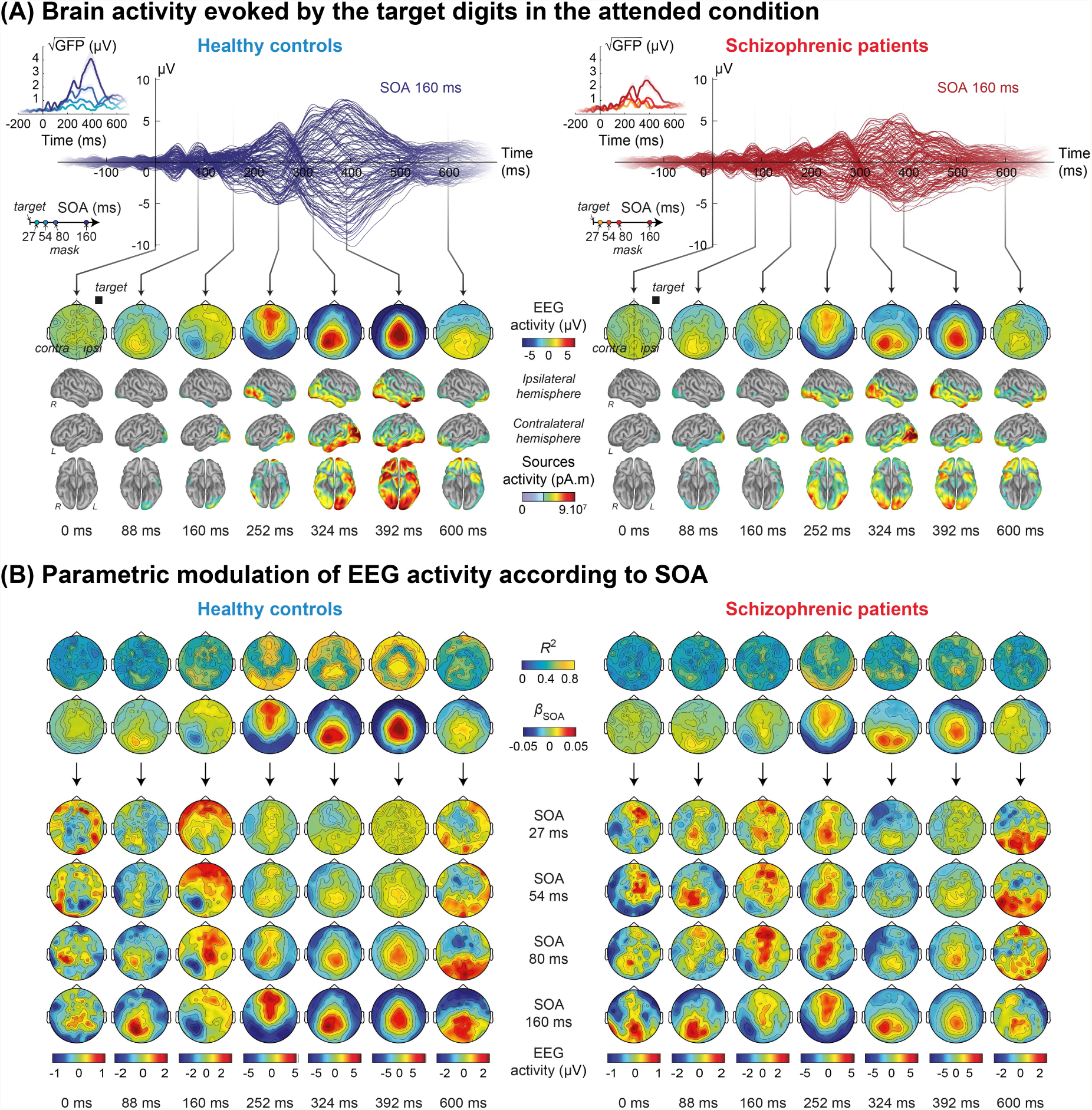
EEG activity evoked by target digits in the attended condition. (A) Time course of brain activity at the longest SOA (i.e. 160 ms) for controls (blue curves on the left) and patients (red curves on the right). Global field potentials are shown in inset as a function of time and SOA. Specific time points were selected, corresponding topographies and source reconstructions are presented below, providing an overview of brain activity evoked by the target as a function of time. Shaded area around the curve represents one standard error of the mean. (B) Topographical maps of both explained variance (R^2^) and regression coefficient (Β) from a linear regression of EEG signals’ amplitude on SOA, performed at each electrode and time point. Below, classical EEG voltage topographies are shown for each time point (horizontally) and for each SOA (vertically).

#### 3.2.1. ERP components amplitudes

In order to examine which of the ERP components evoked by a masked stimulus persist under a condition of inattention, we first tested whether the amplitude of each component was significantly different from zero at the longest SOA (160 ms) under attended and unattended conditions. In the control group, under the attended condition (see Figure 4A), the amplitude of all ERP components was significantly different from zero (P1: t_15_ = 3.10, p = 0.007; N1: t_15_ = −4.95, p < 0.001; N2: t_15_ = −6.25, p < 0.001; P3: t_15_ = 10.83, p < 0.001), while under unattended conditions (see Figure 4B), only the amplitude of the N1 and N2 components was significantly different from zero (N1: t_15_ = −3.35, p = 0.004; N2: t_15_ = −4.54, p < 0.001; P1: t_15_ = −0.05, p = 0.962, BF = 1/3.9; P3: t_15_ = −0.35, p = 0.732, BF = 1/3.7). Similar results were observed in the patient group under attended condition (P1: t_15_ = 4.31, p < 0.001; N1: t_15_ = −3.70, p = 0.002; N2: t_15_ = −3.70, p = 0.002; P3: t_15_ = 6.31, p < 0.001) but only the N2 amplitude was significantly different from zero under unattended condition (N2: t_15_ = −3.91, p = 0.001; P1: t_15_ = −0.09, p = 0.930, BF = 1/3.9; N1: t_15_ = −0.85, p = 0.408, BF = 1/2.9; P3: t_15_ = −0.49, p = 0.635, BF = 1/3.5). For both groups, the P3 component totally vanished under unattended conditions. The results therefore indicate that unattended stimuli could trigger ERPs up to ^~^270 ms after they were presented, but failed to induce a detectable P3 component.

#### 3.2.2. Group effects

We then explored the group effects, with the hypothesis that late ignition would be reduced in the patient group under attended condition. Factorial ANOVAs were conducted on each target-evoked EEG component, with within-subject factors of SOA (27, 54, 80 and 160 ms) and attention (attended or unattended), a between-subjects factor of group (patients or controls), and subject identity as a random factor. The results are summarized in table 2 and time-course ERP amplitude are shown in Figure 4.

P3 was the only component for which a significant overall difference between schizophrenic patients and healthy controls was observed. For the P3, group also significantly interacted with SOA across all attention conditions (F_3,90_ = 6.47, p < 0.001) and the triple interaction group × SOA × attention was significant (F_3,90_ = 6.41, p < 0.001, see Table 2, model 1).

To further explore this group difference, we conducted an ANOVA on the P3 component in each SOA condition, with factors of attention (attended or unattended) and group (patients or controls) and subject as a random factor. It revealed a significant group effect for long SOAs (80 ms: F_1,30_ = 5.80, p = 0.023; 160 ms: F_1,30_ = 5.20, p = 0.030) and a significant interaction between group and attention for SOA 160 ms only (F_1,30_ = 4.74, p = 0.037).

A Group × SOA effect on P3 was observed under attended conditions but not under unattended conditions (attended, see Model 2A: group × SOA: F_3,90_ = 8.53, p < 0.001; unattended, see Model 2U: group × SOA: F_3,90_ = 0.95, p = 0.421, BF = 1/8.0). No main effect of group was observed for P3 either in the attended (see Model 2A: F_1,30_ = 1.65, p = 0.209, BF = 1/2.1) or in the unattended condition (see Model 2U: F_1,30_ = 0.17, p = 0.683, 1/BF =4.3). T-test, however, confirmed a significant difference between patients and controls for P3 under attended conditions at the longest SOAs (SOA 80 ms: Welch t_29.3_ = 2.10, p = 0.044; SOA 160 ms: t_29.6_ = 2.50, p = 0.018, see figure 4A).

For the earlier ERP components P1, N1 and N2, no significant group effect or interaction was observed (see detailed statistics in Table 2, models 1, 2A and 2U).

To sum up, the main impairment observed in schizophrenic patients was an abnormal P3 for long SOAs under attended condition. The significant group × SOA interaction suggested an abnormal ignition at long SOAs. The significant group × attention interaction for the longest SOA suggested that this effect was restricted to the attended condition.

#### 3.2.3. SOA effects

We then turned to the effects of SOA to explore how ERP amplitudes were modulated by the available time to process the target before the mask disrupted it. Across groups and conditions, SOA had a significant main effect on N1, N2 and P3 (Model 1: N1: F_3,90_ = 21.88, p < 0.001; N2: F_3,90_ = 35.01, p < 0.001; P3: F_3,90_ = 45.35, p < 0.001) but not for P1 (F_3,90_ = 1.64, p = 0.187, BF = 1/18.0).

The modulation of ERP amplitude by SOA under attended condition is shown in Figure 3B and 4A. Results from controls (Table 2, model 3AC) replicated previous findings (Del Cul et al., 2007). P1 was not significantly affected by masking (SOA effect: F_3,45_ = 2.26, p = 0.094, BF = 1/1.6). On the contrary, N1, N2 and P3 amplitudes significantly increased with SOA (N1: F_3,45_ = 12.74, N2: F_3,45_ = 29.49, P3: F_3,45_ = 69.58, p < 0.001, *R*^2^ larger than 0.4 for both components, see Figure 3B).

Similarly, in the patient group, there was a significant effect of SOA on N1, N2 and P3 (N1: F_3,45_ = 6.60, N2: F_3,45_ = 13.42, P3: F_3,45_ = 16.82, p < 0.001, see Table 2, model 3AP). The significant effect of SOA on P1 amplitude vanished when excluding SOA = 160 ms (F_2,30_ = 1.47, p = 0.247, BF = 1/3.1). As mentioned above (see Group effects section), the only significant interaction that was observed between group and SOA occurs for the P3, reflecting a much reduced effect of SOA on P3 amplitude in patients compared to controls (F_1,105_ = 6.33, p < 0.001). Such a reduced modulation of P3 by SOA in patients may underpin their lower objective and subjective behavioural performances compared to controls in the attended task (see Discussion).

In the unattended condition, in both groups, SOA had a significant effect on N1 and N2 (see Table 2, model 3UC and 3UP) but not on P1 and P3. The significant increase in N1 and N2 suggested that sensory information could still be processed as a function of SOA even when unattended (see Discussion). These SOA effects did not differ between patients and controls under unattended conditions (see Table 2, model 2U).

To sum up, SOA had an effect on N1 and N2 in both attended and unattended conditions without any significant difference between groups, and on P3 under attended conditions only, with a significant difference between patients and controls.

#### 3.2.4. Attention effects and interactions between attention and SOA

We now report the interactions involving the attentional manipulation to see which component is significantly amplified by attention. Across groups and SOA, attention had a significant effect on all ERP components (P1: F_1,30_ = 4.92, p = 0.034; N1: F_1,30_ = 13.14, p = 0.001; N2: F_1,30_ = 5.14, p = 0.031; P3: F_1,30_ = 69.06, p < 0.001, see Table 2, model 1) and a significant interaction between SOA and attention was observed for all ERP components (P1: F_3,90_ = 4.04, p = 0.010; N1: F_3,90_ = 3.60, p = 0.017; N2: F_3,90_ = 12.01, p < 0.001; P3: F_3,90_ = 67.11, p < 0.001), compatible with the idea that attention modulates the rate of accumulation of sensory information per unit of time (see Discussion).

No significant interaction between group and attention was observed (P1: F_1,30_ = 0.06, p = 0.810, BF = 1/5.0; N1: F_1,30_ = 1.17, p = 0.288, BF = 1/2.7; N2: F_1,30_ = 0.002, p = 0.961, BF = 1/5.3; P3: F_1,30_ = 0.68, p = 0.415, BF = 1/3.3). The triple interaction between group, SOA and attention did not reach significance for the early components (P1: F_3,90_ = 0.45, p = 0.716, BF = 1/9.4; N1: F_3,90_ = 1.20, p = 0.314, BF = 1/11.8; N2: F_3,90_ = 1.76, p = 0.160, BF = 1/6.8), but did for the P3 (F_3,90_ = 6.41, p < 0.001). Indeed, the attentional modulation effect on P3 was lower in the patients compared to the controls (see Table 2, model 4C and 4P; controls: F_3,45_ = 77.43, p < 0.001; patients: F_3,45_ = 13.09, p < 0.001: F_3,45_ = 13.09, p < 0.001) and this difference was significant for the longest SOA (group × attention for SOA 160 ms: F_1,30_ = 4.74, p = 0.037, see Group effect section).

No significant difference between patients and controls was observed for N1. However, in the control group, a main effect of attention and an interaction SOA × attention were significant for N1 (attention: F_1,15_ = 17.70, p < 0.001; SOA × attention: F_3,45_ = 3.41, p = 0.025, see Table 2, model 4C) while it was not the case in the patient group (attention: F_1,15_ = 2.35 p = 0.146, BF = 1.2; SOA × attention: F_3,45_ = 1.79, p = 0.163, BF = 1/4.6, see Table 3 model 4P).

To sum up, across groups, an attentional modulation was observed for all components and had a significant interaction with SOA. This effect of attention was different between the two groups for the P3 at the longest SOA.

**Table 2.** *F, p*-values and Bayes factors from ANOVAs on ERP components.

| ERP component | P1 | N1 | N2 | P3 |
| --- | --- | --- | --- | --- |
| **Model 1: Amplitude ~ Group x SOA x Attention** |  |  |  |  |
| **Group** | $F_{1,30} = 0.06$<br>$p = 0.803$<br>$BF = 1/6.7$ | $F_{1,30} = 2.03$<br>$p = 0.165$<br>$BF = 2.5$ | $F_{1,30} = 0.24$<br>$p = 0.627$<br>$BF = 1/5.3$ | $F_{1,30} = 1.67$<br>$p = 0.207$<br>$BF = 1/3.2$ |
| **SOA** | $F_{3,90} = 1.64$<br>$p = 0.187$<br>$BF = 1/18.0$ | $F_{3,90} = 21.88$<br>**$p < 0.001$** | $F_{3,90} = 35.01$<br>**$p < 0.001$** | $F_{3,90} = 45.35$<br>**$p < 0.001$** |
| **Attention** | $F_{1,30} = 4.92$<br>**$p = 0.034$** | $F_{1,30} = 13.14$<br>**$p = 0.001$** | $F_{1,30} = 5.14$<br>**$p = 0.031$** | $F_{1,30} = 69.05$<br>**$p < 0.001$** |
| **Group x SOA** | $F_{3,90} = 0.55$<br>$p = 0.649$<br>$BF = 1/17.9$ | $F_{3,90} = 1.12$<br>$p = 0.347$<br>$BF = 1/14.5$ | $F_{3,90} = 0.01$<br>$p = 0.961$<br>$BF = 1/23.3$ | $F_{3,90} = 6.47$<br>**$p < 0.001$** |
| **Group x attention** | $F_{1,30} = 0.06$<br>$p = 0.810$<br>$BF = 1/5.0$ | $F_{1,30} = 1.17$<br>$p = 0.288$<br>$BF = 1/2.7$ | $F_{1,30} = 0.00$<br>$p = 0.961$<br>$BF = 1/5.3$ | $F_{1,30} = 0.68$<br>$p = 0.415$<br>$BF = 1/3.3$ |
| **SOA x attention** | $F_{3,90} = 4.04$<br>**$p = 0.010$** | $F_{3,90} = 3.60$<br>**$p = 0.017$** | $F_{3,90} = 12.01$<br>**$p < 0.001$** | $F_{3,90} = 67.11$<br>**$p < 0.001$** |
| **Group x SOA x attention** | $F_{3,90} = 1.64$<br>$p = 0.716$<br>$BF = 1/9.4$ | $F_{3,90} = 1.20$<br>$p = 0.314$<br>$BF = 1/11.8$ | $F_{3,90} = 1.76$<br>$p = 0.160$<br>$BF = 1/6.8$ | $F_{3,90} = 6.41$<br>**$p < 0.001$** |
| **Model 2A: Amplitude ~ Group x SOA under attended conditions** |  |  |  |  |
| **Group effect** | $F_{1,30} = 0.20$<br>$p = 0.658$<br>$BF = 1/4.8$ | $F_{1,30} = 2.57$<br>$p = 0.119$<br>$BF = 2.5$ | $F_{1,30} = 0.09$<br>$p = 0.769$<br>$BF = 1/4.8$ | $F_{1,30} = 1.65$<br>$p = 0.209$<br>$BF = 1/2.1$ |
| **SOA effect** | $F_{3,90} = 4.38$<br>**$p = 0.006$** | $F_{3,90} = 18.14$<br>**$p < 0.001$** | $F_{3,90} = 38.89$<br>**$p < 0.001$** | $F_{3,90} = 74.04$<br>**$p < 0.001$** |
| **Group x SOA** | $F_{3,90} = 0.80$<br>$p = 0.498$<br>$BF = 1/6.8$ | $F_{3,90} = 0.91$<br>$p = 0.442$<br>$BF = 1/7.7$ | $F_{3,90} = 0.82$<br>$p = 0.486$<br>$BF = 1/8.8$ | $F_{3,90} = 8.53$<br>**$p < 0.001$** |
| **Model 2U: Amplitude ~ Group x SOA under unattended conditions** |  |  |  |  |
| **Group effect** | $F_{1,30} = 0.00$<br>$p = 0.983$<br>$BF = 1/5.3$ | $F_{1,30} = 0.50$<br>$p = 0.487$<br>$BF = 1/3.1$ | $F_{1,30} = 0.35$<br>$p = 0.557$<br>$BF = 1/3.8$ | $F_{1,30} = 0.17$<br>$p = 0.683$<br>$1/BF = 4.3$ |
| **SOA effect** | $F_{3,90} = 0.56$<br>$p = 0.644$<br>$BF = 1/18.9$ | $F_{3,90} = 5.62$<br>**$p = 0.001$** | $F_{3,90} = 9.47$<br>**$p < 0.001$** | $F_{3,90} = 0.54$<br>$p = 0.655$<br>$BF = 1/18.0$ |
| **Group x SOA** | $F_{3,90} = 0.13$<br>$p = 0.940$<br>$BF = 1/11.3$ | $F_{3,90} = 1.52$<br>$p = 0.216$<br>$BF = 1/6.5$ | $F_{3,90} = 0.49$<br>$p = 0.687$<br>$BF = 1/8.9$ | $F_{3,90} = 0.95$<br>$p = 0.421$<br>$BF = 1/8.0$ |
| **Model 3AC: Amplitude ~ SOA for controls under attended conditions** |  |  |  |  |
| **SOA effect** | $F_{3,45} = 2.26$<br>$p = 0.094$<br>$BF = 1/1.6$ | $F_{3,45} = 12.74$<br>**$p < 0.001$** | $F_{3,45} = 29.49$<br>**$p < 0.001$** | $F_{3,45} = 69.58$<br>**$p < 0.001$** |

| Model 3AP: Amplitude ~ SOA for patients under attended conditions |  |  |  |  |
| --- | --- | --- | --- | --- |
| SOA effect | $F_{3,45} = 2.86$<br>$p = 0.047$ | $F_{3,45} = 6.60$<br>$p < 0.001$ | $F_{3,45} = 13.42$<br>$p < 0.001$ | $F_{3,45} = 16.82$<br>$p < 0.001$ |
| Model 3UC: Amplitude ~ SOA for controls under unattended conditions |  |  |  |  |
| SOA effect | $F_{3,45} = 0.44$<br>$p = 0.724$<br>$BF = 1/10.1$ | $F_{3,45} = 4.43$<br>$p = 0.008$ | $F_{3,45} = 4.05$<br>$p = 0.013$ | $F_{3,45} = 1.41$<br>$p = 0.252$<br>$BF = 1/6.1$ |
| Model 3UP: Amplitude ~ SOA for patients under unattended conditions |  |  |  |  |
| SOA effect | $F_{3,45} = 0.30$<br>$p = 0.822$<br>$BF = 1/10.5$ | $F_{3,45} = 3.06$<br>$p = 0.038$ | $F_{3,45} = 5.61$<br>$p = 0.002$ | $F_{3,45} = 0.41$<br>$p = 0.746$<br>$BF = 1/9.8$ |
| Model 4C: Amplitude ~ Attention x SOA in control group |  |  |  |  |
| Attention effect | $F_{1,15} = 1.97$<br>$p = 0.181$<br>$BF = 3.9$ | $F_{1,15} = 17.70$<br>$p < 0.001$ | $F_{1,15} = 3.71$<br>$p = 0.073$<br>$BF = 2.0$ | $F_{1,15} = 34.43$<br>$p < 0.001$ |
| SOA effect | $F_{3,45} = 1.64$<br>$p = 0.193$<br>$BF = 1/8.9$ | $F_{3,45} = 14.51$<br>$p < 0.001$ | $F_{3,45} = 22.84$<br>$p < 0.001$ | $F_{3,45} = 43.63$<br>$p < 0.001$ |
| Attention x SOA | $F_{3,45} = 1.65$<br>$p = 0.191$<br>$BF = 1/4.3$ | $F_{3,45} = 3.41$<br>$p = 0.025$ | $F_{3,45} = 12.42$<br>$p < 0.001$ | $F_{3,45} = 77.43$<br>$p < 0.001$ |
| Model 4P: Amplitude ~ Attention x SOA in patient group |  |  |  |  |
| Attention effect | $F_{1,15} = 3.01$<br>$p = 0.103$<br>$BF = 1.9$ | $F_{1,15} = 2.35$<br>$p = 0.146$<br>$BF = 1.2$ | $F_{1,15} = 2.01$<br>$p = 0.177$<br>$BF = 1/1.1$ | $F_{1,15} = 35.54$<br>$p < 0.001$ |
| SOA effect | $F_{3,45} = 0.83$<br>$p = 0.487$<br>$BF = 1/14.6$ | $F_{3,45} = 8.65$<br>$p < 0.001$ | $F_{3,45} = 14.09$<br>$p < 0.001$ | $F_{3,45} = 10.42$<br>$p < 0.001$ |
| Attention x SOA | $F_{3,45} = 2.75$<br>$p = 0.054$<br>$BF = 1/4.4$ | $F_{3,45} = 1.79$<br>$p = 0.163$<br>$BF = 1/4.6$ | $F_{3,45} = 3.03$<br>$p = 0.039$ | $F_{3,45} = 13.09$<br>$p < 0.001$ |

**Figure 4.**
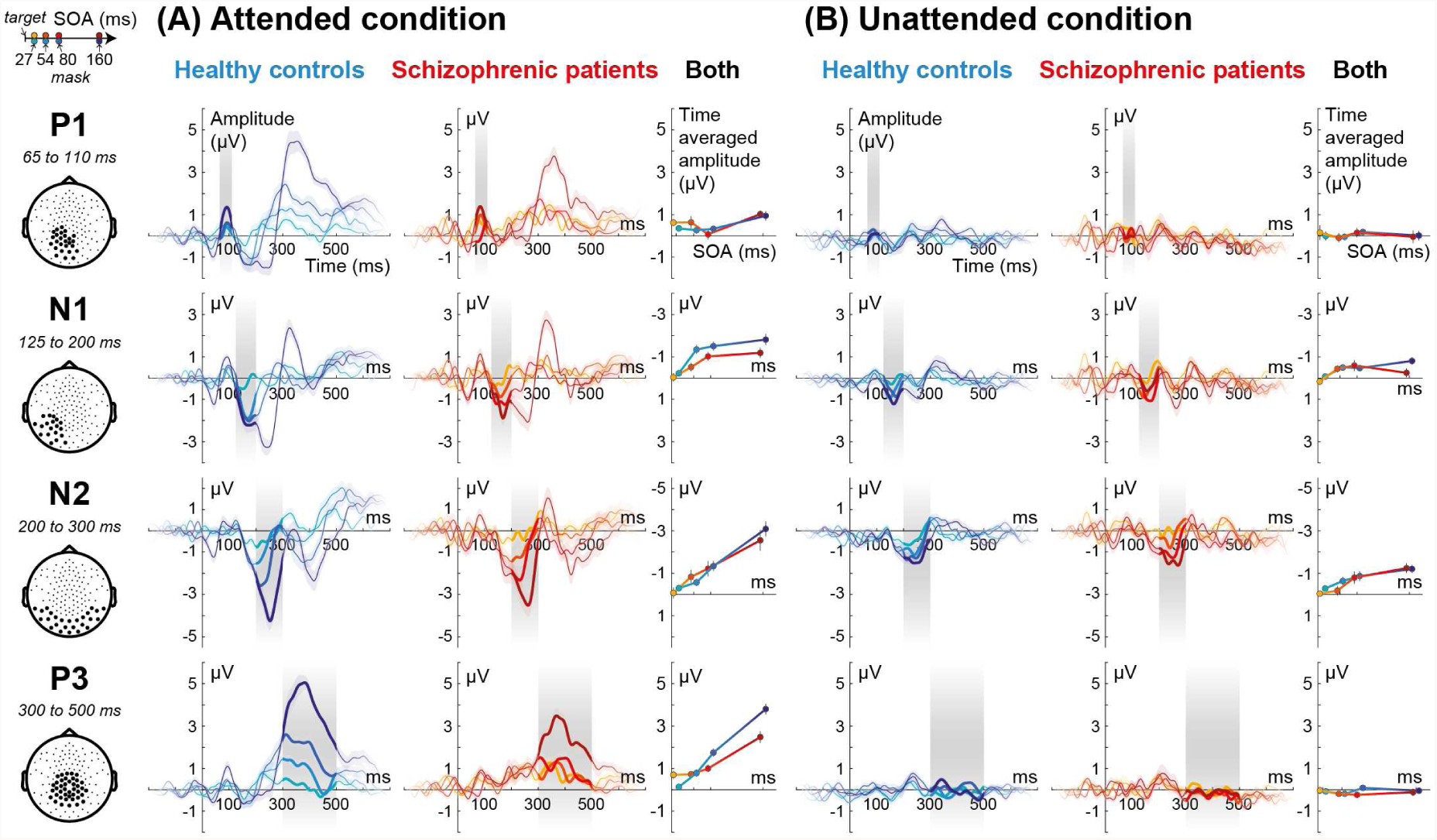
Modulation of ERP components as a function of SOA. Each subplot shows the time course of ERPs as a function of SOA in the control and the patient groups under attended and unattended conditions. For each component, the preselected cluster of electrodes is depicted by black dots in the topographies at left. Preselected time-windows of interest, used for statistical analysis, are shown by grey rectangles. Colored shaded area around the curves represents one standard error of the mean. The averaged amplitude of each component in this window is also plotted (column marked “both”). Error bars represent one standard error of the mean.

## 4. Discussion

### 4.1 Summary of the results

We measured the effect of top-down attention on visual stimuli whose degree of masking varied by modulating the target-mask SOA duration. Our main results can be summarized as follows.

First, in the healthy control group, when subjects attended to the masked target, we replicated our previous observations of a monotonic increase of ERPs’ amplitude (N1, N2, P3) as the target-mask interval increased (Del Cul et al., 2007). Inattention reduced the amplitude of all ERP components, decreased the slope with which the N1 and N2 varied as a function of SOA, and led to a complete disappearance of the P3 component. Attention therefore had both a modulatory influence on early perceptual processing and an all-or-none effect on the late P3 component.

Second, no difference was observed between the schizophrenic patient and the healthy control groups under unattended condition. In particular, the modulation of cerebral activity by SOA took place normally for N1 and N2. However, patients’ consciousness thresholds, as assessed by subjective visibility and objective performance were abnormally elevated, and their P3 component was reduced relative to controls in the attended condition for long SOAs. Earlier components (P1, N1, N2) were not significantly affected.

### 4.2 Persistence of bottom-up processing under unattended condition

One of the main goals of our experiment was to examine which of the ERP components evoked by a masked stimulus persist under a condition of inattention. The unattended condition, which involved continuous attention to the color of the fixation point, was specifically designed to induce a complete withdrawal of spatial, temporal and executive attention resources to the peripheral masked stimulus. For several minutes, this peripheral stimulus was therefore completely task-irrelevant and ignored. As a consequence, we could not record any behavioural or introspective measurements as to how this stimulus was processed. An indirect indication of strong inattention, however, was that target presence and target-mask SOA had no effect on the performance of the color estimation task, although this performance was far from ceiling.

We predicted that, in spite of this strong inattention, peripheral stimuli should still elicit early visual ERP components, up to about 300 ms, but should no longer yield a P3 waveform. This pattern is exactly what was observed. Under the unattended condition, the P1 component was strongly attenuated. The N1 and N2 components, although attenuated as well, were still observable and reflected a clear activation of occipito-temporal cortices similar to what was observed under attended condition. Furthermore, both N1 and N2 components continued to be modulated by SOA, suggesting that the accumulation of perceptual evidence from the target digit continued to occur even without attention. The results were however different for the P3, which collapsed to an undetectable level. These results are compatible with our previous postulate that brain states prior to 300 ms post-target (i.e. P1, N1 and N2) correspond to a series of largely automatic “pre-conscious” perceptual stages (Dehaene et al., 2006), while latter ones such as the P3 reflects an all-or-none stage of conscious access (Dehaene and Changeux, 2011; Del Cul et al., 2007). Source reconstruction also suggests that the brain correlates of conscious access are reflected by a highly distributed set of activations involving the bilateral inferior frontal, anterior temporal and inferior parietal cortices. On the contrary, when attention is distracted during the inattention task, we observe a spatially reduced brain activity that was restricted to posterior visual and occipital areas. A relative preservation of early activations (P1, N1, N2) was previously described under other inattention paradigms such as the attentional blink (Harris et al., 2013; Marti et al., 2012; Sergent et al., 2005; Vogel and Luck, 2002) or inattentional blindness (Pitts et al., 2011). Such a preservation of early brain processes may explain why priming effects are repeatedly observed both in inattentional blindness and attentional blink conditions.

### 4.3 Attention and the amplification of evidence accumulation

The original contribution of the present experimental paradigm is to demonstrate, through the manipulation of SOA, that attention amplifies sensory evidence and its accumulation rate relative to strong inattention. The literature on attention has primarily focused on the issues of whether attention modulates early as well as late processes. Our study confirms that attention can have a strong modulating influence on early components, although withdrawal of attention does not completely eradicate them (Feng et al., 2012; Hillyard and Anllo-Vento, 1998; Kastner and Ungerleider, 2000; Luck and Ford, 1998; Woodman and Luck, 2003; Zotto and Pegna, 2015). However, our study points to another way in which attention impacts perceptual processing. By manipulating the SOA between the target and a subsequent mask, we found that many processing stages integrate stimulus information, in the sense that their activation increases monotonically with SOA. This was particularly the case for N2 which, as noted earlier (Del Cul et al., 2007), starts at a fixed delay relative to target onset, ends at a fixed delay relative to mask onset, and appears to increase linearly in amplitude as a function of the interval elapsed between these two events. These three properties suggest that N2 might reflect an accumulation of sensory evidence that continues until it is interrupted by the mask. Moreover, the present results extend these findings by showing that the slope of the SOA modulation, i.e. the amount of integrated information per unit of time, also called “drift rate”, can be modulated by attention. Under conditions of inattention, the modulation of ERP amplitude by SOA was indeed either weakened or simply entirely absent, suggesting that attention might impact the temporal integration constant of perceptual networks. Crucially, the target was presented for the same duration in all conditions (14 ms). It therefore seems that the brain buffers this sensory information while being able to accumulate samples from it through a series of processing stages, with a slope proportional to attention, until another concurrent information (i.e. the mask) reinitializes the sensory buffer, thereby stopping the accumulation process. In summary, top-down attention seems to enable a specific mode of amplification and integration in which a fixed quantity of sensory evidence provided at input is able to trigger a series of successive stages of increasingly amplified activation, and which ultimately translates into a global ignition.

In accordance with previous theoretical models, we propose that peripheral brain processors accumulate sensory information which will be consciously perceived if it crosses a threshold and accesses a distributed global workspace able to stabilize and make it available to a variety of processes (Baars, 1993; Dehaene, 2011; Dehaene and Changeux, 2011; Lafuente and Romo, 2006). Importantly, accumulation of evidence can be carried out on unconscious perceptual information (Vlassova et al., 2014; Vorberg et al., 2003) and may precede conscious access (Vorberg et al., 2003). Our results concur with this idea by showing a significant increase in ERP amplitude with SOA even under unattended condition. However, they also refine these findings, indicating that such unconscious evidence accumulation process can be amplified by top-down attention and suggesting that conscious perception corresponds to a threshold crossing in evidence accumulation (Dehaene, 2011; Kang et al., 2017; King and Dehaene, 2014; Ploran et al., 2007; Shadlen and Kiani, 2011).

### 4.4 P3 increases beyond the minimal consciousness threshold

Prior research, using different criteria, indicates that the presence or absence of a P3 component tightly correlates with conscious access (using a variety of paradigms with fixed stimuli and variable subjective experience: Babiloni et al., 2006; Del Cul et al., 2007; Fernandez-Duque et al., 2003; Lamy et al., 2008; Pins and Ffytche, 2003; Sergent et al., 2005). Recently, this view has been challenged by concurrent hypotheses proposing that P3 might reflect post-perceptual processing rather than truly being a neural correlate of consciousness. Indeed, P3 was observed to be absent even for consciously perceivable stimuli when these were task-irrelevant (Pitts et al. 2011,2014; Shafto and Pitts, 2015).

In our previous work (Del Cul et al., 2007), SOA varied only in the range 16-100 ms. Over this range and under attended condition, we observed a sigmoidal variation of objective and subjective indices of target visibility, and we found that P3 amplitude closely tracked this sigmoidal shape. Here, however, by extending the SOA to longer values (27-160 ms), we observed that the P3 amplitude continued to increase in the range 100-160 ms where subjective visibility reached a fixed ceiling. Still, P3 amplitude again closely tracked visibility in the sense that it was nil at SOA = 27 ms, precisely when subjects reported that stimuli were essentially invisible, and then increased for larger SOAs when the stimuli became visible. The P3 thus showed a threshold-like non-linearity at short SOAs (see Figure 4A), unlike other waveforms such as the N2 which was already observable for stimuli that were judged invisible (i.e. SOA = 27 ms).

Such a continued P3 increase at long SOAs was unexpected and indicates a departure for the close parallelism previously suggested between conscious reports and P3 size (Babiloni et al., 2006; Del Cul et al., 2007; Fernandez-Duque et al., 2003; Lamy et al., 2008; Pins and Ffytche, 2003; Sergent et al., 2005). This aspect of our results suggests that, like previous ERP stages, P3 may reflect an evidence-accumulation process, but within a high-level cognitive route associated with subjective experience and reportability, above and beyond the mere sensori-motor mapping level (Dehaene, 2011; Del Cul et al., 2009; King and Dehaene, 2014; Shadlen and Kiani, 2011). Several other studies have indeed shown how P3 is associated with the formation of decisions and can reflect evidence accumulation (Gold and Shadlen, 2007; O’Connell et al., 2012; Twomey et al., 2015) as well as post-decision confidence (Boldt and Yeung, 2015; Murphy et al., 2015). Given those studies, it seems possible that the binary subjective measure that we have used (seen/unseen) did not fully do justice to the rich introspection that subjects had about target visibility. Had we measured a more continuous parameter such as confidence or clarity, it seems possible that one or several of such behavioural indices would have grown continuously with SOA, paralleling the observed increase in P3 size.

### 4.5 Abnormal attentional amplification in schizophrenia

Behaviourally, we replicated the previous findings according to which schizophrenic patients suffer from a higher objective and subjective thresholds for conscious perception during masking (Butler et al., 2003; Charles et al., 2017; Dehaene et al., 2003a; Del Cul et al., 2006; Green et al., 1999,2011; Herzog and Brand, 2015; Plomp et al., 2013). The main goal of our study was to evaluate whether this deficit was associated with impairments of bottom-up and/or top-down processing. The results were clear-cut: schizophrenic patients, compared to healthy controls, showed anomalies in evoked brain activity only under attended conditions for long SOAs: the late non-linear ignition component associated with the P3 component was reduced. However, no difference was found under unattended condition. We emphasize the need for caution in interpreting those null findings in the unattended condition, as they might be due to a lack of power arising from the small sample size (16 patients and 16 controls). Nevertheless, our data were sensitive enough to detect a preservation of the modulation of the N1 and N2 by SOA in the patient group under unattended conditions. In other words, both the target processing and the initial accumulation of evidence as well as its modulation by SOA took place normally in patients when the stimulus was unattended. We therefore conclude that patients’ deficit in perceiving masked stimuli probably mostly arises from a lack of appropriate top-down attentional amplification rather than from a mere bottom-up impairment.

At the level of the P3, the difference between patients and controls was significant only for long SOAs. The patients exhibited a detectable P3 in the attended compared to the unattended condition (see Figure 4) but there was almost no modulation of its amplitude by SOA when SOA was shorted than 80 ms (see Figure 4A). These results are consistent with the behavioural results showing reduced objective performances in the patient group only at long SOAs (Figure 2).

In our work, no significant difference between patients and controls was observed for the N1. This finding contrasts with several previous studies that found a reduced N1 amplitude in the auditory modality (Brockhaus-Dumke et al., 2008; Turetsky et al., 2008) and in several visual masking paradigms (Neuhaus et al., 2011; Wynn et al., 2013). Careful examination of the present results suggests that a non-significant difference in N1 amplitude may be observable in Figure 4A for SOA > 27 ms. Moreover, N1 topography also seems to be different in patients and controls at SOA 160 ms (see Figure 3). According to source reconstruction, posterior negative cerebral activity is more ventral and more bilateral in patients compared to controls at SOA 160 ms (see Sources in Figure 3A). For SOA 54 and 80 ms, N1 is still visible in controls but not in patients and a frontal positivity is present in controls but not in patients for SOA 27 and 54 ms (Figure 3B). Because of our small sample size (n = 16 in each group), we may simply lack enough statistical power to demonstrate a significant statistical difference between groups for N1 under attended conditions, and this effect should be re-investigated in future experiments.

Interestingly, patients showed essentially normal attentional amplification of the P1 and N2 components This result is in line with previous studies suggesting that attentional selection could be preserved when guided by strong bottom-up salience (Gold et al., 2017).

As reviewed in the introduction, some authors proposed that the elevated threshold for conscious access in schizophrenia was due to a specific dysfunction of the magnocellular pathway, while the parvocellular visual pathway was thought to be preserved (Butler et al., 2005,2007; Javitt, 2009; Kim et al., 2006; Martínez et al., 2012). Tapia and Breitmeyer (2011), however, revisited this issue and proposed that magnocellular channels contribute to conscious object vision mainly through a top-down modulation of re-entrant activity in the ventral object-recognition stream. The link between magnocellular circuits and visual masking in schizophrenia was also contested recently, as there seems to be no clear evidence of either hyper or hypo-activity of the magnocellular pathway in schizophrenia (Herzog and Brand, 2015).

If the elevated threshold for conscious perception in schizophrenia was solely due to abnormal bottom-up sensory processing, one would expect subliminal and unattended processing to be abnormal too. However, first, even subtle measures of subliminal priming have repeatedly been shown to be fully preserved in schizophrenia (Dehaene et al., 2003a; Del Cul et al., 2006; for a review, see: Berkovitch et al., 2017) and our results are compatible with these observations since no difference was observed for short SOAs. Second, the present results extend this logic by showed that, following the total withdrawal of spatial, temporal and executive attention, the remaining brain activity evoked by a flashed stimulus is indistinguishable between patients and controls. By hypothesis, this activity should provide a proper measure of bottom-up processing, which therefore appears to be essentially intact.

Consequently, we suggest that previous reports of elevated masking threshold and abnormal conscious processing in schizophrenia (Butler et al., 2003; Charles et al., 2017; Dehaene et al., 2003a; Del Cul et al., 2006; Green et al., 1999; Herzog et al., 2004; Plomp et al., 2013) might stem from late impairments in processing stages associated with the P3 and which, in turn, are associated with the inability to deploy top-down attention. An abnormal P3 and ignition deficits had already been reported in schizophrenia in attended conditions (Bramon et al., 2004; Charles et al., 2017; Jeon and Polich, 2003; Oribe et al., 2015; Qiu et al., 2014) and several studies showed that the difference in cerebral activity between attended and unattended conditions was reduced in schizophrenia (Force et al., 2008; Martínez et al., 2012; Michie et al., 1990). Moreover, other studies suggested impairments in top-down processing (Dima et al., 2010; Plomp et al., 2013) and selective attention (Fuller et al., 2006; Luck et al., 2006) in which schizophrenic patients were characterized by a narrower spotlight of spatial attention termed hyperfocusing (Hahn et al., 2012; Leonard et al., 2017; Sawaki et al., 2017).

The present study therefore corroborates the hypothesis of a top-down impairment in schizophrenic patients and refines previous results by clearly distinguishing bottom-up versus top-down processes and showing that some top-down attentional amplification (underlying P1 and N2 components) can remain preserved in schizophrenia. Tentatively, one may suggest that the activations that were found to be preserved in schizophrenic patients (i.e. P1 and N2, but also P3 for short SOAs) might account for the preservation of subliminal processing.

More broadly, the present results fit with several other physiopathological aspects of schizophrenia (Berkovitch et al., 2017). Schizophrenic patients exhibit anomalies in longdistance anatomical connectivity (Bassett et al., 2008; Benetti et al., 2015; Jones et al., 2006; Kubicki et al., 2005; Sigmundsson et al., 2001) and functional connectivity (Ford et al., 2002; Frith et al., 1995; Lawrie et al., 2002; Vinckier et al., 2014) in distributed networks that are thought to underlie the broadcasting of conscious information in the global workspace (Dehaene and Changeux, 2011). Moreover, the long-range synchrony of gamma and beta-band oscillations is disturbed in schizophrenic patients (Cho et al., 2006; Lee et al., 2003; Mulert et al., 2011; Spencer et al., 2004; Uhlhaas and Singer, 2010), while conscious perception in normal subjects is accompanied by late increases in gamma-band power (Doesburg et al., 2009; Gaillard et al., 2009; Melloni et al., 2007; Wyart and Tallon-Baudry, 2009) and beta-band phase synchrony (Gaillard et al., 2009; Gross et al., 2004; King et al., 2013). Finally, abnormal regulation of NMDA receptors was suggested as a putative core pathology in schizophrenia (Coyle, 2006; Jentsch and Roth, 1999; Olney and Farber, 1995; Stephan et al., 2009). NMDA receptors are broadly involved in connectivity and synaptic plasticity (Stephan et al., 2009) as well as inter-areal synchrony (Rivolta et al., 2015; Uhlhaas and Singer, 2014; van Kerkoerle et al., 2014). Recently, they have been shown to play a specific role in top-down cortico-cortical connectivity and the late amplification of sensory signals (Herrero et al., 2013; Moran et al., 2015; Self et al., 2012; van Loon et al., 2016). In particular, NMDA-receptor antagonists leave intact the feedforward propagation of visual information, and selectively impact on late recurrent processing (Self et al., 2012). NMDA receptor dysfunction could therefore be a plausible cause for the anomaly in conscious perception observed in the present work.

### 4.6 Conclusion

Our study aimed to disentangle how sensory information processing is modulated by bottom-up (SOA) and top-down (attention) factors. We found that, in the absence of attention, bottom-up information was still processed and weakly modulated the early stages of information processing, prior to 300 ms. Attention, however, enabled a strong amplification of sensory signals that, in its late stages, played a decisive part in conscious access. The abnormal consciousness threshold in schizophrenia seems tightly linked to a dysfunction of the latter top-down attentional amplification mechanisms.

## Acknowledgements

This work was supported by INSERM, CEA, Collège de France, Fondation Roger de Spoelberch, and an ERC grant “Neuroconsc” to S.D.; Fondation pour la Recherche Médicale to L.B. and S.D.; a “Frontières du Vivant” doctoral fellowship involving the Ministère de I’Enseignement Superiéur et de la Recherche and Fondation Bettencourt to M.M. We gratefully acknowledge the hospitality of Marion Leboyer.

